# From silence to song: Testosterone triggers extensive transcriptional changes in the female canary HVC

**DOI:** 10.1101/2022.06.13.495861

**Authors:** Meng-Ching Ko, Carolina Frankl-Vilches, Antje Bakker, Nina Sohnius-Wilhelmi, Pepe Alcami, Manfred Gahr

**Affiliations:** Department of Behavioural Neurobiology, Max Planck Institute for Biological Intelligence, Seewiesen, 82319, Germany; Division of Neurobiology, Faculty of Biology, Ludwig-Maximilians-University Munich, Grosshaderner Strasse 2, 82152 Planegg-Martinsried, Germany

## Abstract

Seasonal song production in canaries, influenced by gonadal hormones, is a well-documented phenomenon. We explored testosterone-induced song development in adult female canaries—a behavior rarely exhibited naturally. Gene regulatory networks in the song-controlling brain area HVC were compared at multiple time points (1 hour to 14 days) post-treatment with those of placebo-treated controls, paralleling HVC and song development. Females began vocalizing within four days of testosterone treatment, with song complexity and HVC volume increasing progressively over two weeks. Rapid transcriptional changes involving 2,739 genes preceded song initiation. Over two weeks, 9,913 genes—approximately 64% of the canary’s protein-coding genome—were differentially expressed, with 98% being transiently regulated. These genes are linked to various biological functions, with early changes at the cellular level and later changes affecting the nervous system level after prolonged hormone exposure. Our findings suggest that testosterone-induced song development is accompanied by extensive and dynamic transcriptional changes in the HVC, implicating widespread neuronal involvement. The data reveal extensive transcriptomic changes, including alterations in steroid receptor expression and numerous transcription factors, coinciding with significant neural transformations. These changes underpin the gradual emergence of singing behavior, providing insights into the neural basis of seasonal behavioral patterns.

## Introduction

Adult socio-sexual behaviors in vertebrates are critically regulated by gonadal hormones, particularly androgens and estrogens ^1–3^, which fluctuate in response to environmental conditions ^4,5^. These hormones can induce significant changes in brain sensitivity and structure, leading to altered neural circuit configurations that underpin seasonal behaviors. A prominent example of this phenomenon is the seasonal singing of some songbird species, which coincides with morphological changes in song-controlling brain regions ^6–10^.

Androgens and estrogens exert their effects by binding to their respective receptors, initiating transcription of gene cascades or activating intracellular second messenger pathways ^11,12^. While these hormones may affect several regions of a neural circuit, the impacted gene cascades within a specific brain region seem limited in adults ^13,14^. Unraveling the complex interplay between hormones, gene expression, brain structure, and behavior is particularly challenging when investigating behaviors that emerge slowly over time, such as the seasonal singing of canaries ^15,16^.

Singing behavior in songbirds is a complex process involving intensive integration of sensory inputs and motor outputs ^17^. In many North temperate species, sexually motivated singing is typically limited to males and is sensitive to testosterone and its metabolites ^18,19^. The neural song control system of songbirds ^17,20^ is a target of testosterone due to the abundant expression of androgen and estrogen receptors ^21,22^.

The premotor forebrain nucleus HVC, a sensorimotor integration center of the song system, controls the temporal pattern of singing ^23,24^. As the only song-control nucleus expressing both androgen and estrogen receptors ^25,26^, the HVC is likely subject to extensive hormonal regulation at various organizational levels. Testosterone has been shown to induce angiogenesis in adult female canaries’ HVC within a week ^27^, increase the recruitment of new neurons ^28^, and promote the growth of HVC volume ^29,30^.

Female canaries (*Serinus canaria*) provide a unique model for studying testosterone-induced changes in singing behavior. Typically non-singers, these females possess the necessary underlying circuitry that can be activated by testosterone, leading to the production of male-like songs over several weeks ^31,32^. This model allows for the examination of transcriptional cascades in parallel with the differentiation of the song control system and the progression of song development, without the confounding impact of fluctuating testosterone levels seen in males.

In this study, we investigate the short- and long-term responses of HVC gene expression following testosterone treatment in adult female canaries. We administered testosterone over six specific time periods: 1 hour (h) [T1h], 3 h [T3h], 8 h [T8h], 3 days (d) [T3d], 7 d [T7d], and 14 d [T14d] (Fig. 1A-B and Table 1). Throughout these periods, we monitored vocal activity, gene expression, and neuroanatomical changes. Contrary to our expectation of gradual gene recruitment, our results reveal an immediate and significant impact on HVC gene expression with dynamic patterns over time, affecting 9,913 genes over two weeks, including 843 transcription factors.

**Fig. 1.**
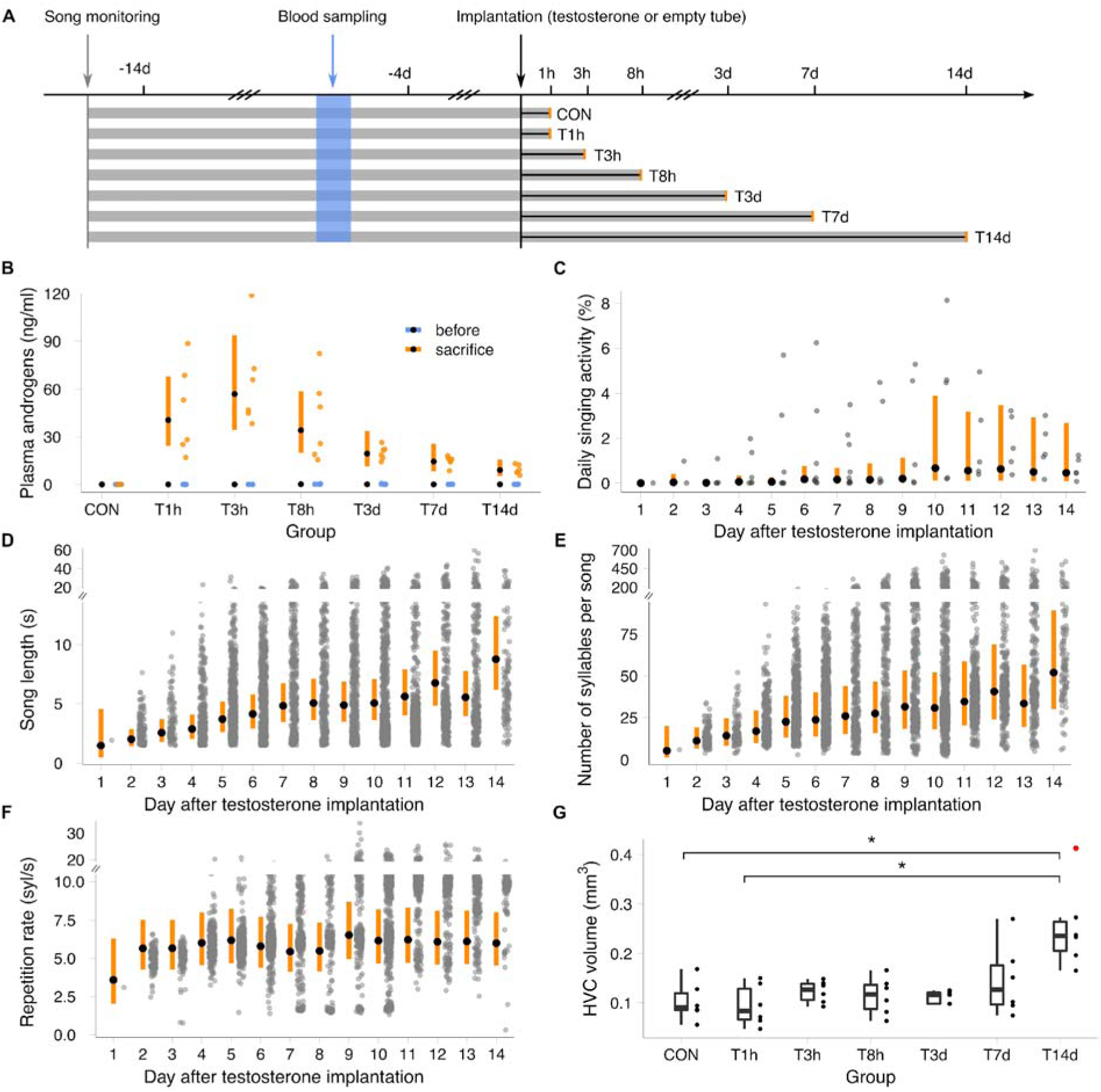
Testosterone implantation affects blood androgen levels, HVC morphology, and song syntax in adult female canaries. A, Experimental timeline. Blood samples were collected before implantation (blue) and at sacrifice (orange). CON: control birds with empty implants; T: testosterone-treated birds for 1 hour (T1h) to 14 days (T14d). B, testosterone significantly increased plasma androgen levels at all time points. Blue: pre-implantation; Orange: post-implantation. Black dots: predicted estimates from linear mixed-effects models; Orange bars: 95% credibility intervals (CrI). See Supplementary Tables 2 and Supplementary Table 3 for the values of the linear mixed-effects models and posterior probabilities. C, daily singing activity (% of 9-hour period) in T7d and T14d groups increased significantly on days 2 and 10 (see also Figure 1 - Figure supplement 3). Each grey dot is one bird. D, song length (seconds, s) in T7d and T14d groups increased daily until day 7, with further increase on day 11. E, number of syllables per song in T7d and T14d groups increased daily post-implantation. F, syllable repetition rate (syllables/second) in T7d and T14d groups increased significantly on days 2 and 9. In D-F, each gray dot represents a measurement originating from one song; black dots show predicted estimates from linear mixed-effects models; orange bars indicate 95% CrI. See Supplementary Table 2 and Supplementary Table 3 for the estimates of the linear mixed-effects models and the posterior probabilities. G, HVC volumes of T14d were significantly different from the control and T1h groups. Boxes: 25th/50th/75th percentiles; Whiskers: values within 1.5 times IQR; Red dots: outliers. *: Holm-adjusted P-value <0.05 (Kruskal-Wallis test followed by Dunn’s post hoc test). Sample sizes of panel B to G are listed in Table 1.

**Table 1.**
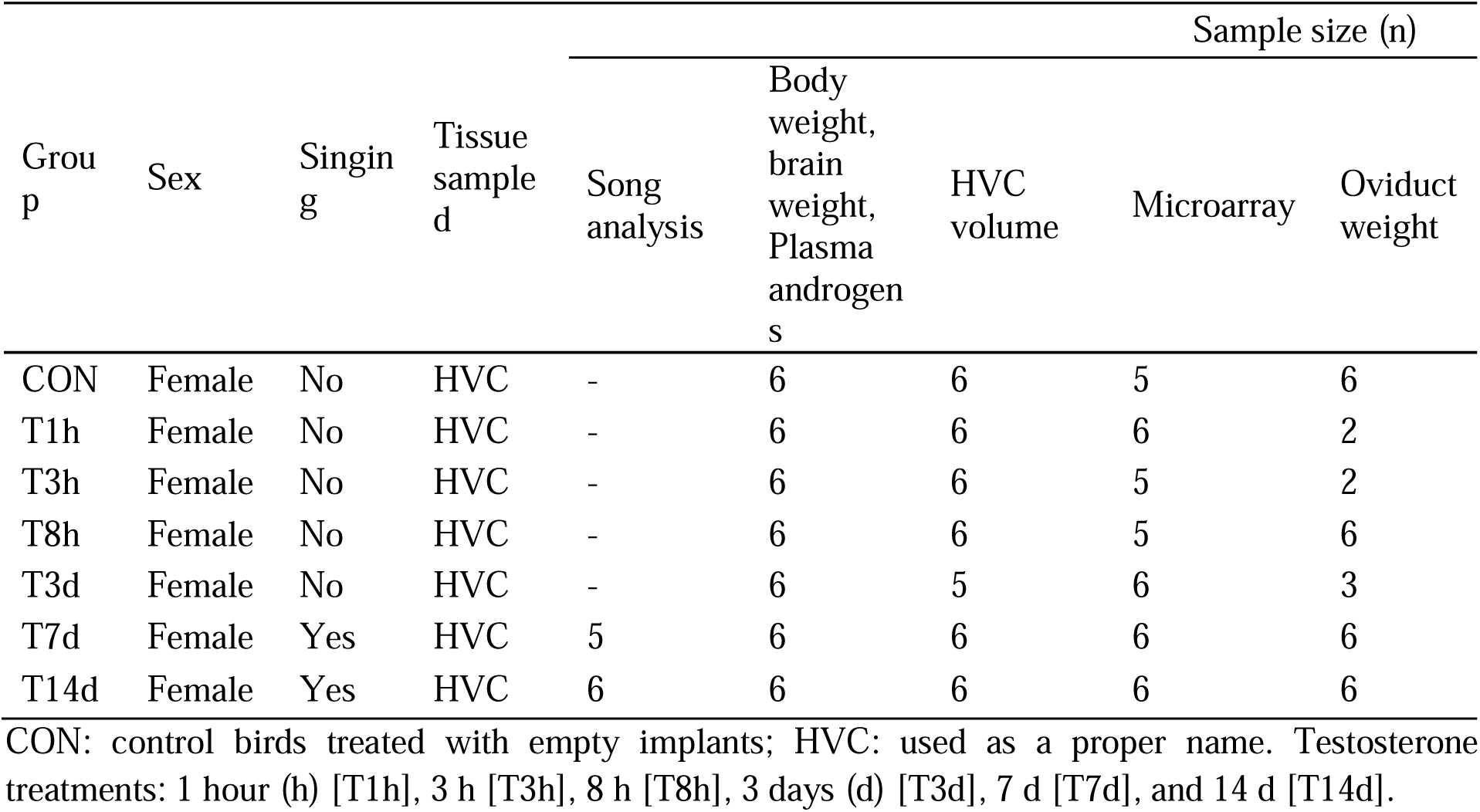
Experimental groups and sample sizes.

## Results

### Testosterone-induced song development

We selected non-singing females to establish a homogeneous baseline for testosterone-induced song development. Song activity was monitored post-hormone implantation to determine the timing of testosterone-induced vocalization (Fig. 1).

Birds in groups T1h, T3h, T8h, and T3d either called or remained silent post-implantation, while five of six birds in group T7d and all birds in group T14d sang (Figure 1 - Figure supplement 1 and Supplementary Table 1). The first recognizable subsongs, characterized as unstructured, low-amplitude vocalizations consisting of three or more syllables, were uttered 4.1 ± 1.6 (mean ± SD) days after implantation. This timing aligns with previous reports in female canaries ^32^ and is comparable to observations in testosterone-implanted male canaries during the non-breeding season ^33^. Notably, one female in the T14 group sang just one day post-treatment. Singing activity increased significantly after days 2 and 10 of testosterone treatment, with no statistical difference in singing activity between days 2 and 9 (Fig. 1C, Supplementary Table 2, and Supplementary Table 3).

### Progressive song development

Early-stage songs displayed substantial inter- and intra-individual variability (Figure 1 - Figure supplement 2 and Supplementary Table 1). Testosterone significantly increased song lengths daily until day 7 and again from day 11 onwards (Fig. 1D, Supplementary Table 2, and Supplementary Table 3). Similarly, the number of syllables per song increased daily post-implantation (Fig. 1E).

Testosterone implantation significantly increased the syllable repetition rate in two distinct steps: initially on day 2 and subsequently on day 9 (Fig. 1F). The maximum observed syllable repetition rate was 26 Hz, consistent with previous findings ^34^.

### Testosterone increased HVC volume after two weeks

HVC volumes were significantly larger in T14d birds compared to controls (Fig. 1G, CON: 0.103 ± 0.040 mm³; T14d: 0.253 ± 0.087 mm³, mean ± SD; Kruskal-Wallis followed by Dunn post hoc test, χ² = 16.15, df = 6, Holm-adjusted P-value = 0.013). HVC volumes in T7d birds (0.146 ± 0.073 mm³) did not significantly differ from controls (Holm-adjusted P-value = 1) or T14d birds (Holm-adjusted P-value = 0.69), likely due to high variability in the T7d group.

Testosterone treatment did not significantly affect body weight, brain weight, or oviduct weight (Figure 1 - Figure supplement 4A-C). HVC volumes normalized to brain weight yielded similar results as non-normalized volumes (Kruskal-Wallis test, χ² = 17.32, df = 6, P-value = 0.0082, Figure 1 - Figure supplement 4D).

These findings align with previous studies demonstrating that systemic testosterone administration leads to detectable HVCs in female canaries ^28–31,34,35^.

### Testosterone dramatically alters HVC transcriptomes

To investigate testosterone’s effect on gene regulation in the HVC, we performed microarray analyses on the six testosterone-treated groups and controls. Testosterone’s impact on transcription was evident as early as 1 hour post-implantation (T1h, Fig. 2A), coinciding with significantly elevated plasma androgen levels (Fig. 1; see Methods). Given our moderate sample sizes, we conducted a power analysis for each differentially expressed gene, considering only genes with power ≥ 0.8 (Figure 2 - Figure Supplement 1).

**Fig. 2.**
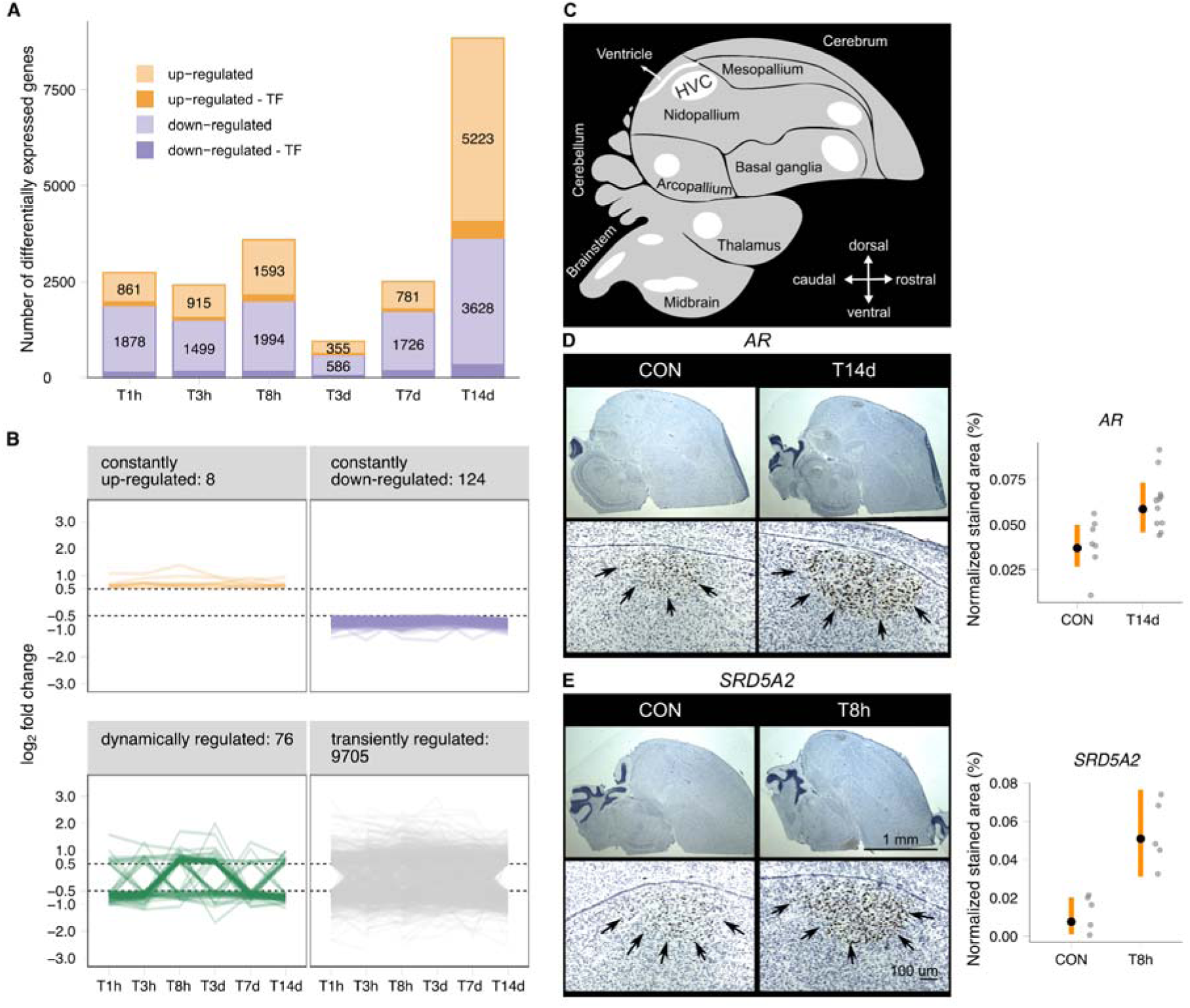
Testosterone rapidly and persistently altered gene expression patterns in the HVC. A, number of differentially expressed genes in the HVC of testosterone-treated birds compared to controls. Darker shades indicate transcription factors (TF). Within one hour (T1h), over 2,700 genes were differentially regulated, with a substantial increase at 14 days (T14d). B, classification of differentially expressed genes based on temporal regulation patterns: 1) constantly upregulated, 2) constantly downregulated, 3) dynamically regulated, and 4) transiently regulated genes (see Supplementary Table 5). C, schematic of songbird brain with song control areas (white) including the HVC. D and E, testosterone increased expression of androgen receptor (*AR*) and 5α-reductase (*SRD5A2*) mRNA in the HVC. Representative RNAScope *in situ* hybridization images were shown for *AR* mRNA (D) and *SRD5A2* mRNA (E) in controls (CON) and testosterone-treated birds (T14d and T8h, respectively). Arrowheads indicate increased expression in treated birds. Quantification confirms microarray-derived expression differences (see Fig. 3A, Supplementary Table 2, and Supplementary Table 3). Data represent the proportion of labeled HVC area relative to total HVC area. Gray dots: individual section values; Black dots: predicted estimates from linear mixed-effects models; Orange bars: 95% confidence intervals (CrI). N = 3 birds per group.

At T1h, 2,739 genes showed significant differential expression compared to controls: 861 upregulated and 1,878 downregulated, including 118 upregulated and 151 downregulated transcription factors (TFs) (Fig. 2A, Figure 2 – Figure supplement 1A, and Supplementary Table 4). The number of differentially expressed genes fluctuated during testosterone treatment, increasing dramatically after 14 days (T14d, Fig. 2A). At T14d, 5,223 genes were upregulated and 3,628 downregulated, including 446 upregulated and 322 downregulated TFs. These 8,851 genes represent 57% of all known protein-coding genes (15,609) in the canary genome ^36^. Across all time points, testosterone treatment affected the expression of 9,913 genes, corresponding to approximately 64% of all canary protein-coding genes.

### The vast majority of testosterone-induced genes are only transiently affected

To understand the temporal dynamics of testosterone-altered gene expression, we classified differentially expressed genes into four categories: 1) constantly upregulated (at all six time points), 2) constantly downregulated, 3) dynamically regulated (direction of regulation varied between time points), and 4) transiently regulated (altered expression at least at one but not all time points) (Fig. 2B). Contrary to our expectation of a gradual increase in affected genes over time, we observed a rapid and massive impact characterized by a dynamic, nonlinear pattern. The majority of differentially expressed genes (9,705) were transiently regulated, while only 208 genes fell into the constant-regulated classes combined (Fig. 2B). Further analysis of the transient category revealed that 45% (4,338) of these genes were differentially regulated at just one time point (Figure 2 - Figure Supplement 1C).

Constantly upregulated genes included *KIF5C* (kinesin Family Member 5C) and *NCAM1* (neural cell adhesion molecule 1). *KIF5C* mutations are associated with cortical malformations and microcephaly ^37^, while *NCAM1* is involved in various brain processes, including neurite growth, axonal and dendritic elongation, and neuronal migration ^38^ (Supplementary Table 5).

Among the constantly downregulated genes, several were associated with synapse organization and function, including *PSEN1* (presenilin 1), *TSC2* (tuberous sclerosis complex 2), *VPS35* (vacuolar protein sorting 35), *CAMK2G* (calcium/calmodulin dependent protein kinase II gamma), and the serotonin receptor *HTR2C* (5-hydroxytryptamine receptor 2C) (Supplementary Table 5).

### Testosterone induces dynamic changes in steroidogenesis, hormone metabolism, and vascularization genes

Microarray results revealed significant changes in genes involved in steroidogenesis, hormone metabolism, and vascularization, with distinct temporal dynamics observed across various time points post-testosterone implantation (Fig. 3A).

**Fig. 3.**
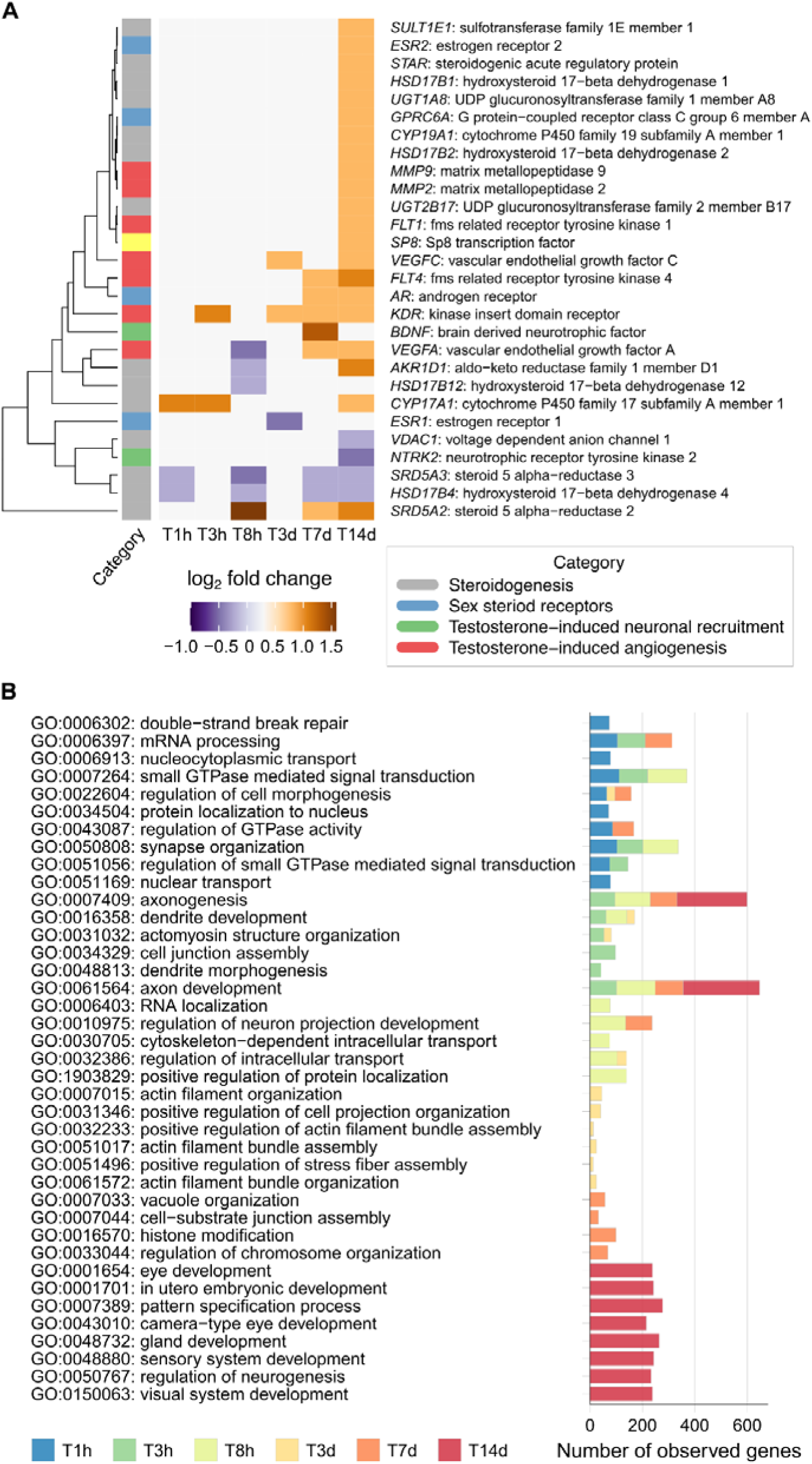
Temporal dynamics of testosterone-induced gene regulation in the HVC. A, dendrogram showing expression levels of selected gene groups during testosterone treatment compared to control HVCs (log_2_ fold change). Groups include gonadal steroid receptor genes (blue), steroidogenesis genes (gray), testosterone-induced angiogenesis genes (red), and neuronal recruitment genes (green). See Figure 3 - Figure Supplement 1 for bird-level normalized expression levels of *AR*, *SRD5A2*, *ESR1*, and *ESR2*. B, GO term enrichment analysis of biological processes involving differentially expressed genes during progressive testosterone action on the HVC. The top ten most significant GO terms (based on Benjamini-Hochberg P-values) are displayed for each time point. The stacked bar graph shows a cumulative number of genes observed across time points (see Supplementary Table 6 for full results). Biological processes at cellular levels occur early (1 hour), while organ/system level processes emerge later (14 days).

Steroidogenesis and hormone metabolism genes showed marked changes. At the early time point of 1 hour, *CYP17A1* (cytochrome P450 17A1) was upregulated, while *HSD17B4* (hydroxysteroid 17-beta dehydrogenase 4) and *SRD5A3* (steroid 5 alpha-reductase 3) were downregulated. By 8 hours, *SRD5A2* (5α-reductase type 2) was upregulated, while *AKR1D1* (aldo-keto reductase family 1 member D1) and *HSD17B12* (hydroxysteroid 17-beta dehydrogenase 12) were downregulated. At 14 days, *CYP17A1*, *CYP19A1* (aromatase), *SRD5A2*, *AKR1D1*, *UGT1A8*, and *UGT2B17* were upregulated.

Steroid receptors also exhibited significant changes. The androgen receptor (*AR*) was upregulated at 7 and 14 days, indicating increased androgen sensitivity. Estrogen receptors showed dynamic regulation, with *ESR1* (estrogen receptor 1) downregulated at 3 days and *ESR2* (estrogen receptor 2) upregulated at 14 days. *GPRC6A*, a potential membrane androgen receptor, was upregulated at 14 days.

Vascularization-related genes also showed significant changes. *KDR* (kinase insert domain receptor) was upregulated at multiple time points from 3 hours to 14 days (except T8h), while *VEGFC* (vascular endothelial growth factor C) showed upregulation at later time points. Other vascularization genes, including *FLT4*, *FLT1*, and *VEGFA*, exhibited dynamic regulation patterns across the treatment period.

These changes in mRNA expression highlight testosterone’s impact on the neural and vascular environment in the HVC. The upregulation of genes involved in steroidogenesis and vascularization suggests that testosterone promotes both hormonal activity and increased vascular support, potentially enhancing neural function and plasticity. The dynamic regulation of androgen and estrogen receptors further underscores the complexity of hormone-driven gene regulation in the brain. Further studies are necessary to confirm these findings and elucidate the mechanisms underlying these gene expression changes.

### Testosterone-induced genes are first associated with cellular changes and later with neural system changes

Gene Ontology (GO) term enrichment analysis was performed to predict potential biological processes affected by testosterone implantation in the HVC at different time points (Fig. 3B and Supplementary Table 6). One hour after testosterone treatment, genes associated with processes such as synapse organization (GO:0050808) and regulation of cell morphogenesis (GO:0022604) were enriched. By three hours, genes related to neural projections, including axon development (GO:0061564) and dendrite development (GO:0016358), showed enrichment. Many processes associated with cellular anatomical changes demonstrated continuous enrichment from T1h to T14d. KEGG pathway enrichment analysis yielded similar findings (Figure 3 - Figure supplement 2A and Supplementary Table 7).

At T14d, the biological processes of differentially expressed genes shifted dramatically. Alongside earlier active processes associated with cellular differentiation, broader functions such as organ formation and system development emerged, including in utero embryonic development (GO:0001701) and sensory system development (GO:0048880). Additional GO term enrichment analysis of 3,269 genes differentially regulated only at T14d also revealed mainly system-level processes (Figure 3 - Figure supplement 2B and Supplementary Table 8). These observations suggest significant changes throughout the entire HVC after approximately 14 days of testosterone treatment.

These findings indicate that testosterone treatment initially influences cellular and morphological processes in the HVC, progressively leading to broader systemic and neural developmental changes over time, culminating in significant alterations after 14 days. This temporal progression underscores the complex interplay between hormonal regulation and gene expression in orchestrating structural and functional transformations in the HVC.

## Discussion

Our study reveals unprecedented transcriptomic plasticity in the adult brain, demonstrating how hormonal signals can rapidly and extensively remodel neural circuits underlying complex learned behaviors (Fig. 4). By examining testosterone-induced song development in female canaries, we uncovered a dynamic gene regulatory landscape in the HVC, a key song control nucleus.

**Fig. 4.**
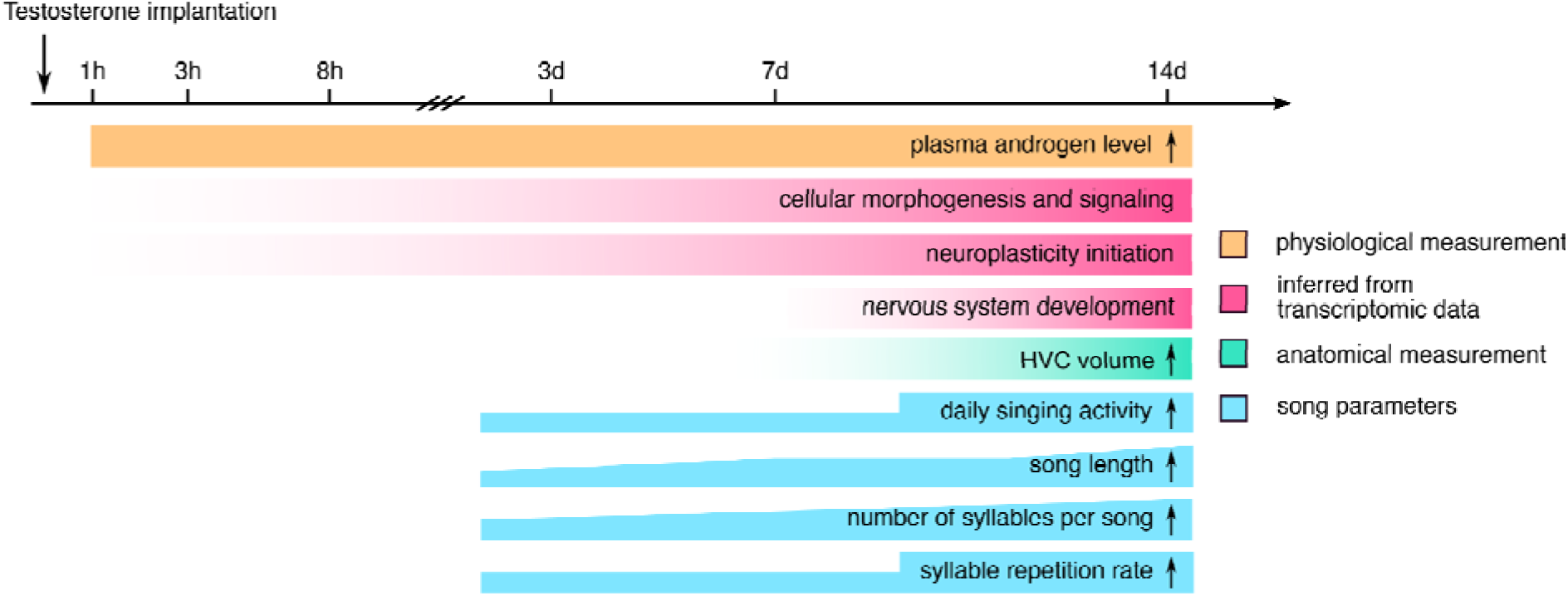
Timeline of HVC differentiation and song development following testosterone treatment. This figure illustrates the temporal progression of HVC organizational levels derived from transcriptome data, HVC morphology measured as volume, and song activity and syntax measurements after testosterone treatment. The intensities of colors indicate the abundance or prominence of features. Biological processes related to the circulatory system and neuron development precede those associated with the nervous system development of HVC and the emergence of song. The appearance of canary-typical song patterns coincides with the system development of HVC. HVC morphology (volume) reaches maturity only after approximately 2 weeks of treatment, coinciding with the most differentiated song patterns uttered by the canaries. This timeline provides a comprehensive view of the parallel developments in gene expression, neuroanatomy, and behavior throughout the testosterone-induced song-learning process.

The temporal dynamics of gene expression in the HVC following testosterone treatment were striking. Within one hour of testosterone exposure, 2,739 genes exhibited differential expression in the HVC, preceding any observable changes in song behavior. This number escalated to 8,851 genes by day 14, coinciding with the emergence of male-like song patterns. Remarkably, over 98% of the 9,913 differentially expressed genes were regulated at specific time points, suggesting that distinct developmental stages require unique transcriptional networks.

The temporal specificity of gene regulation closely mirrored the progression of song development. Early transcriptional changes (T1h to T3d) are primarily related to neuronal and cellular differentiation, potentially priming the HVC for initial vocalizations. As birds advanced to producing plastic songs around T7d, we observed enrichment in genes associated with neuronal differentiation and BDNF signaling, known to regulate the recruitment and survival of adult-born neurons ^39^. The most extensive transcriptional changes occurred at T14d, correlating with significant HVC enlargement and the production of near-crystallized songs.

The dramatic shift in gene expression at T14d is particularly noteworthy. Alongside earlier active processes associated with cellular differentiation, broader functions such as organ formation and system development emerged. This suggests that testosterone-induced song development involves a restructuring of the entire HVC, reminiscent of embryonic developmental processes. This parallel between adult plasticity and embryonic development provides intriguing insights into the mechanisms of hormone-induced behavioral changes in adults.

The scale of gene regulation observed in this study far exceeds previous reports in CNS literature, which typically identify fewer than 2,000 differentially expressed genes ^13,14,40–50^. This discrepancy may be attributed to the high expression of androgen and estrogen receptors in HVC neurons ^21,22,51,52^ and the cellular diversity of the HVC ^53^. However, we cannot exclude the possibility that species-specific factors, methodological variations, and the extended duration of our study also contributed to these differences. Moreover, the striking parallels between late-stage (T14d) gene expression patterns and those observed in embryonic organ formation suggest that testosterone may trigger adult behaviors through extensive transcriptomic restructuring of relevant brain areas.

Our findings raise intriguing questions about the mechanisms underlying this remarkable neural plasticity. The rapid onset of transcriptional changes suggests direct regulation by androgen and estrogen receptors, while the subsequent waves of gene expression likely involve complex cascades of secondary effectors. The extensive remodeling of the HVC transcriptome may reflect not only changes in neuronal gene expression but also alterations in glial and vascular components ^27–29,31,32^, contributing to the observed increase in HVC volume.

While our study focused on the HVC, singing involves a broader song control system ^17^. Future research should investigate testosterone’s effects on other brain areas within this system to provide a more comprehensive understanding of the neural basis of singing behavior. Additionally, disentangling the direct effects of testosterone from those induced by increased singing activity presents an important challenge for future studies, as singing itself can influence gene expression in the HVC ^54^.

In conclusion, our results demonstrate that adult song development involves a series of precisely timed, large-scale transcriptional events that reshape the entire HVC. This work provides unprecedented insights into the molecular underpinnings of hormone-induced neural plasticity and lays the groundwork for future studies exploring the relationship between gene regulation and complex behavioral outputs in the adult brain. Furthermore, it highlights the remarkable ability of the adult brain to undergo extensive, orchestrated changes in response to hormonal signals, reminiscent of developmental processes typically associated with embryonic stages.

## Materials and Methods

### Animals

Forty-two adult female canaries (*Serinus canaria*, at least one-year-old) were used in this study (Table 1). Birds were bred in the animal facility of the Max Planck Institute for Biological Intelligence, Seewiesen, Germany. Housing and welfare adhered to the European Directives for the protection of animals used for scientific purposes (2010/63/EU), with protocols approved by the Government of Upper Bavaria (AZ-No. 55.2/54-2531-68/12). The study followed ARRIVE guidelines ^55^, with a checklist provided in the manuscript.

Bird sex was confirmed by PCR using primers P2 and P8 for the CHD genes ^56^ and by post-mortem inspection of the female reproductive system. Food and water were provided *ad libitum*. Six to eight weeks before the experiment (starting no earlier than September), the light schedule was gradually adjusted to short-day conditions (light:dark = 9:15 hours). After acclimation to this cycle, singing was monitored. Birds were divided into a control group and six experimental groups (Table 1). Experimental groups received testosterone implants for varying durations: 1 hour (T1h), 3 hours (T3h), 8 hours (T8h), 3 days (T3d), 7 days (T7d), and 14 days (T14d) (see Method: Testosterone Implantation).

### Song monitoring

About 5% of female canaries sing spontaneously without hormone manipulation ^57^. To ensure that vocal activity in our experiment was indeed triggered by testosterone implantation, birds used in this study were housed alone in sound-attenuated boxes (70 × 50 × 50 cm) and all vocalizations were recorded continuously for two weeks before testosterone implantation and until killing. The microphone (TC20, Earthworks, Milford, New Hampshire) in each silenced box was connected to a PR8E amplifier (SM Pro Audio, Melbourne, Australia) that fed into an Edirol USB audio recording device (Edirol UA 1000, Roland, Los Angeles, California) connected to a computer. Vocalizations were recorded at a sampling rate of 44.1 kHz and a resolution of 16 bits using Sound Analysis Pro 2011 software ^58^.

### Song analysis

Song data analysis was limited to the T7d and T14d groups, as testosterone-treated birds began singing unstable subsongs on average by day 4. Nine hours of recordings were analyzed daily from day 0 until euthanasia (day 7 or 14). Data from T7d and T14d birds were combined for the first 7 days’ analysis, as their songs did not differ significantly (Figure 1 - Figure Supplement 3, Supplementary Table 2, and Supplementary Table 3).

Analysis was performed using Multi_Channel_Analyser (MCA), a custom MATLAB program (version R2016b, MathWorks) with a graphical user interface, as previously described (Ko et al. 2020) and publicly available (https://doi.org/10.5281/zenodo.1489098). MCA generates sound spectrograms using a fast Fourier transform with a sliding time window of 294 samples and 128 sample overlap, resulting in a spectrogram resolution of 150 Hz (frequency) and 3.76 milliseconds (time).

Syllable recognition involved three steps due to background noise. First, sound segments (songs and calls) were manually selected by visual inspection. Second, song segments longer than 2 seconds were selected. Third, syllables within selected segments were automatically detected using MCA with parameters set to: “High-pass filter” 2.0 kHz, “Threshold_wave_amplitude” 0.001, and “Threshold_syllable_freq” 50 Hz. Sounds shorter than 5 milliseconds were removed.

We analyzed four song-level parameters (song length, number of syllables per song, syllable repetition rate, and slope coefficient α ^59–61^) and four syllable-level parameters (syllable length, syllable spacing, peak frequency, and Wiener entropy).

Song length was calculated from the first syllable’s start to the last syllable’s end. Syllable repetition rate was defined as syllables per second. Syllable length and spacing were calculated from syllable timestamps. Peak frequency was the frequency of maximum power in the syllable spectrogram. Wiener entropy, a measure of signal noise, was log-transformed for a broader range (0 represents white noise, infinity represents pure tone) ^58^.

### Testosterone implantation

Custom-made testosterone implants were prepared by packing testosterone (Sigma-Aldrich, 86500, Saint Louis, Missouri) into 7-mm-long SilasticTM tubes (Dow Corning, Midland, Michigan; 1.47 mm inner diameter, 1.96 mm outer diameter, 0.23 mm thickness). Both ends were sealed with silicone elastomer (3140, Dow Corning). Implants were cleaned with 100% ethanol to remove any testosterone particles and tested for leakage by overnight immersion in ethanol; moist implants were discarded. One day before implantation, implants were incubated overnight in 0.1 M phosphate-buffered saline (PBS) to ensure immediate testosterone release upon implantation ^28^.

Subcutaneous implantation began immediately after lights-on at 8:30 am, with a 20-minute interval between each bird to account for euthanasia timing. Control animals received empty 7-mm silicone tubes sealed with silicone elastomer. After the designated implantation period (1h, 3h, 8h, 3d, 7d, or 14d, Table 1), birds were euthanized with an isoflurane overdose. Body weight was recorded, and brains and oviducts were dissected, weighed, frozen on dry ice, and stored at −80□ until further use. Oviducts of eleven birds were not collected or weighed during dissection. At euthanasia, all testosterone implants were present and contained testosterone.

### Radioimmunoassay of plasma androgens

To confirm increased circulating androgen levels following hormone manipulation, we performed a radioimmunoassay on blood samples collected before and after testosterone implantation (Fig. 1A). Approximately 100 μl of blood was collected twice from the wing vein—once around 4 days before implantation and once immediately before euthanasia. Samples were collected between 8 and 11 am within 3 minutes to minimize handling effects ^62^. Blood was centrifuged (5,000 rpm, 10 min) to separate plasma. Testosterone metabolites in plasma were measured using a commercial antiserum to testosterone (T3-125, Endocrine Sciences, Tarzana, CA) as described by ^63^. Standard curves and sample concentrations were calculated with Immunofit 3.0 (Beckman Inc., Fullerton, CA) using a four-parameter logistic curve fit and corrected for individual recoveries.

Testosterone concentrations were assayed in duplicates across four separate assays, with a mean extraction efficiency of 84.0 ± 6.8% (N = 120). The lower detection limits for the assays were 0.32-0.44 pg per tube, and all samples were above this limit. Intra-assay coefficients of variation for a chicken plasma pool were 4.4%, 4.1%, 9.8%, and 2.6%, and the inter-assay coefficient of variation was 4.3%. The testosterone antibody cross-reacts significantly with 5α-dihydrotestosterone (44%), so measurements include a fraction of 5α-DHT and are referred to as plasma androgen levels.

Testosterone implants increased plasma androgen levels in all treated groups (Supplementary Table 2 and Supplementary Table 3). Pre-implantation levels were statistically similar across groups (159 ± 449 pg/ml, mean ± SD). Implantation increased plasma androgen levels by at least 28-fold above baseline (range: 17 to 88 ng/ml) after one hour, with no significant differences between testosterone-treated groups post-implantation (CON 0.0472 ± 0.0240; T1h 46.9 ± 28.2; T3h 53.9 ± 14.8; T8h 35.6 ± 24.0; T3d 20.4 ± 4.47; T7d 14.8 ± 3.29; T14d 9.31 ± 2.94 ng/ml).

### Brain sectioning

Brains were sagittally sectioned using a cryostat (Jung CM3000, Leica, Wetzlar, Germany) in either 40 μm × 4 + 20 μm × 2 (N = 17) or 50 μm × 4 + 14 μm × 3 (N = 33) sections. Thick sections (40 or 50 μm) were mounted on glass slides for microdissection and microarray analysis, while thin sections (20 or 14 μm) were mounted on RNase-free Thermo Scientific™ SuperFrost Plus™ slides (J1800AMNZ, Thermo Fisher Scientific, Waltham, MA) for Nissl staining or RNAScope® *in situ* hybridization. All sections were stored at −80°C until further processing.

### Measurement of the HVC volume

Thin serial sections (20 or 14 μm) were Nissl-stained with 0.1% thionine solution and covered with Roti-Histokitt II embedding medium (Roth). HVC areas were measured using ImageJ2 (Fiji distribution) ^64,65^. All brains were coded to ensure blinded evaluation. Volumes were calculated from summed-area measurements multiplied by section thickness and spacing. The Nissl-stained sections of one T3d bird were of poor quality, so its HVC volume was not measured.

### RNAScope^®^ in situ hybridization assay

We performed RNAScope® *in situ* hybridization for *AR* and *SRD5A2* mRNA expression on 20 or 14 μm thick sections using the RNAScope® 2.0 HD Detection Kit (Advanced Cell Diagnostics, Newark, California), following the manufacturer’s protocols ^66^. Three birds were used per probe at each time point, with three slices per bird. Probe details are summarized in Supplementary Table 9. Stained sections were imaged using a Leica DM6000 B microscope and quantified with ImageJ2. Chromogenic particles in the HVC were measured and normalized to the HVC area. The Color Threshold function (RGB; red: 0-47, green: 0-165, blue: 0-160) and Analyze Particles function (minimum size: 8 μm²) were used to count and measure particle areas. User-defined macro files are available on GitHub (https://github.com/maggieMCKO/TimeLapseTestoFemaleCanary/blob/ce9261d1cf96e760351733657434e499dea35174/RNAscopeQuantification/Marco_quantify_density_singleProbes_quantify_area.ijm).

### Microarray procedures and annotation

Forty-two adult female canaries (six birds per group) were planned for microarray analysis following the method proposed by ^67^. However, two birds were excluded due to insufficient RNA, resulting in thirty-nine canaries being used (see Table 1). The procedure was described in ^68^.

We extracted total RNA by dissecting the HVC from 40 or 50 μm sections under a stereomicroscope, referencing adjacent Nissl-stained sections. The HVC is located ventral to the hippocampus, lateral ventricle, and caudo-dorsal to the lamina mesopallialis in sagittal sections. Approximately 20 slices per HVC were dissected and transferred into an Eppendorf tube with 340 μl of RLT buffer (Qiagen, Valencia, CA). RNA was extracted using the RNeasy® Micro Kit (Qiagen), including a DNase digest step, and assessed for quality using an Agilent 2100 Bioanalyzer and a Nanodrop 1000 spectrometer (Thermo Fisher Scientific). All samples had RNA integrity numbers (RIN) > 7.

Purified RNA samples (≥ 100 ng) were processed and hybridized using the Ambion WT Expression Kit and the Affymetrix WT Terminal Labeling and Controls Kit (Thermo Fisher Scientific). The cDNA was hybridized to the Custom Affymetrix Gene Chip® MPIO-ZF1s520811 Exon Array, validated for cross-species hybridization ^36,69^. Hybridization was performed for 16 hours at 45°C and 60 rpm. Arrays were washed, stained, and scanned using the Affymetrix GeneChip Fluidics Station 450 and GeneChip scanner 3000 7G. CEL files were generated with Affymetrix GeneChip Command Console® Software (AGCC), and hybridization quality was assessed using Affymetrix Expression Console™ software.

The custom array included 5.7 million male zebra finch-specific probes corresponding to 25,816 transcripts. Over 90% of transcripts were annotated to 12,729 human orthologous genes using public (Ensembl, GenBank, UniProt, DAVID) ^70–75^) and commercial databases (v 21 El Dorado, Genomatix, Precigen Bioinformatics Germany GmbH (PBG), Munich, Germany). The microarray data generated in this study have been deposited in NCBI’s Gene Expression Omnibus ^76^ and are accessible through the GEO Series accession number GSE118522.

### Analysis of differentially expressed genes

Differential expression analyses were performed using the R package limma ^77^, following the workflow of Klaus and Reisenauer ^78^. Six pairwise comparisons were conducted using a “Paired Samples” design (limma user guide, section 9.4), comparing each testosterone-treated group to the control group. Comparisons were corrected for multiple testing using the Benjamini-Hochberg method, with significant differentially expressed transcripts defined by an adjusted P-value < 0.05 and a minimum differential expression of |log2(fold change)| ≥ 0.5.

Significantly differentially expressed transcripts were annotated to human orthologous genes. For transcripts from the same gene, average expression was calculated if all transcripts were regulated in the same direction. If both up- and down-regulated transcripts were present, transcripts of the minority direction (<40%) were discarded, and the average expression was calculated from the remaining transcripts (≥60%). Transcripts without human orthologous gene annotation were removed before subsequent analyses.

A power analysis was conducted for each probe-set in the pairwise differential expression analysis due to moderate sample sizes. Effect sizes were calculated using Robust Multichip Average (RMA)-normalized expression from CEL files, and power was estimated using the “pwr.t.test” function (two-sided, α = 0.05) from the “pwr” R package (v1.3-0) ^79^. Genes with at least one probe-set having a power ≥ 0.8 were considered high-power differentially expressed genes. In total, 98% of differentially expressed genes identified by limma with |log2(fold change)| ≥ 0.5 had a power ≥ 0.8.

The differential analysis results align with published data at the overlapping time point (T14d, Fig. 2E), where elevated mRNA levels of vascular endothelial growth factor receptor (KDR) were reported ^27^. RNAScope® in situ hybridization assays for androgen receptor (AR) and 5α-reductase 2 (SRD5A2) confirmed the microarray results (Fig. 2).

### Enrichment analysis of Gene Ontology (GO)-terms and KEGG pathways

We used the ‘enrichGO’ and ‘enrichKEGG’ functions of the clusterProfiler v4.12.0 R package ^80^ to predict the putative biological functions (GO and KEGG) of genes of interest. P values were corrected for multiple comparisons using the Benjamini-Hochberg procedure, and results were plotted using the R package ggplot2 ^81^.

### Experimental design and statistical analysis

All statistical analyses were performed in R ^82^. Linear mixed-effect models analyzed fixed effects using the ‘lme4’ ^83^ and ‘arm’ ^84^ packages in a Bayesian framework with non-informative priors. Gaussian error distribution was assumed and model assumptions were verified by visual inspection of residuals. Plasma androgen levels, daily song rate, song length, and syllable repetition rate were log-transformed; normalized stained HVC areas (RNAScope®) were square root transformed. Estimates were simulated 10,000 times using the ‘sim’ function to extract 95% credible intervals (CrI) for the mean of simulated values, representing estimate uncertainty (Gelman and Hill, 2006). Effects were considered statistically meaningful if the CrI did not include zero or if the posterior probability of the mean difference was higher than 0.95 ^85^. Linear mixed-effect models, model estimates, and posterior probabilities are detailed in Supplementary Table 2 and Supplementary Table 3. Predicted estimates, 2.5% and 97.5% CrI, and raw data were plotted for visualization. Statistically meaningful differences are inferred if one group’s CrI does not overlap with the mean estimate of the other group.

### Materials availability statement

The datasets presented in this study are available in online repositories. Microarray CEL files can be accessed on NCBI’s Gene Expression Omnibus <u>GSE118522</u>. The custom MATLAB sound analysis program, Multi_Channel_Analyser (MCA), is available on GitLab https://doi.org/10.5281/zenodo.1489098. Processed microarray data, plasma androgen levels, HVC volume measurements, and song parameters are deposited on Dryad doi:10.5061/dryad.5hqbzkh8c. All analysis and visualization scripts are available on GitHub https://github.com/maggieMCKO/TimeLapseTestoFemaleCanary.

## Acknowledgments.

We thank Dr. Stefan Leitner for providing breeding canaries, Roswitha Brighton and David Witkowski for maintaining the colony, and Dr. Wolfgang Goymann and Monika Trappschuh for conducting the radioimmunoassay of plasma androgens. We also appreciate the valuable comments on previous versions of this manuscript from Drs. Maude Baldwin, Falk Dittrich, Vincent Van Meir, and Michiel Vellema. Additionally, we acknowledge the support and training provided by the International Max Planck Research School for Organismal Biology.

## Competing interests

The authors declare no competing financial interests.

## Supplementary Figures and Legends

**Figure 1 – Figure supplement 1.**
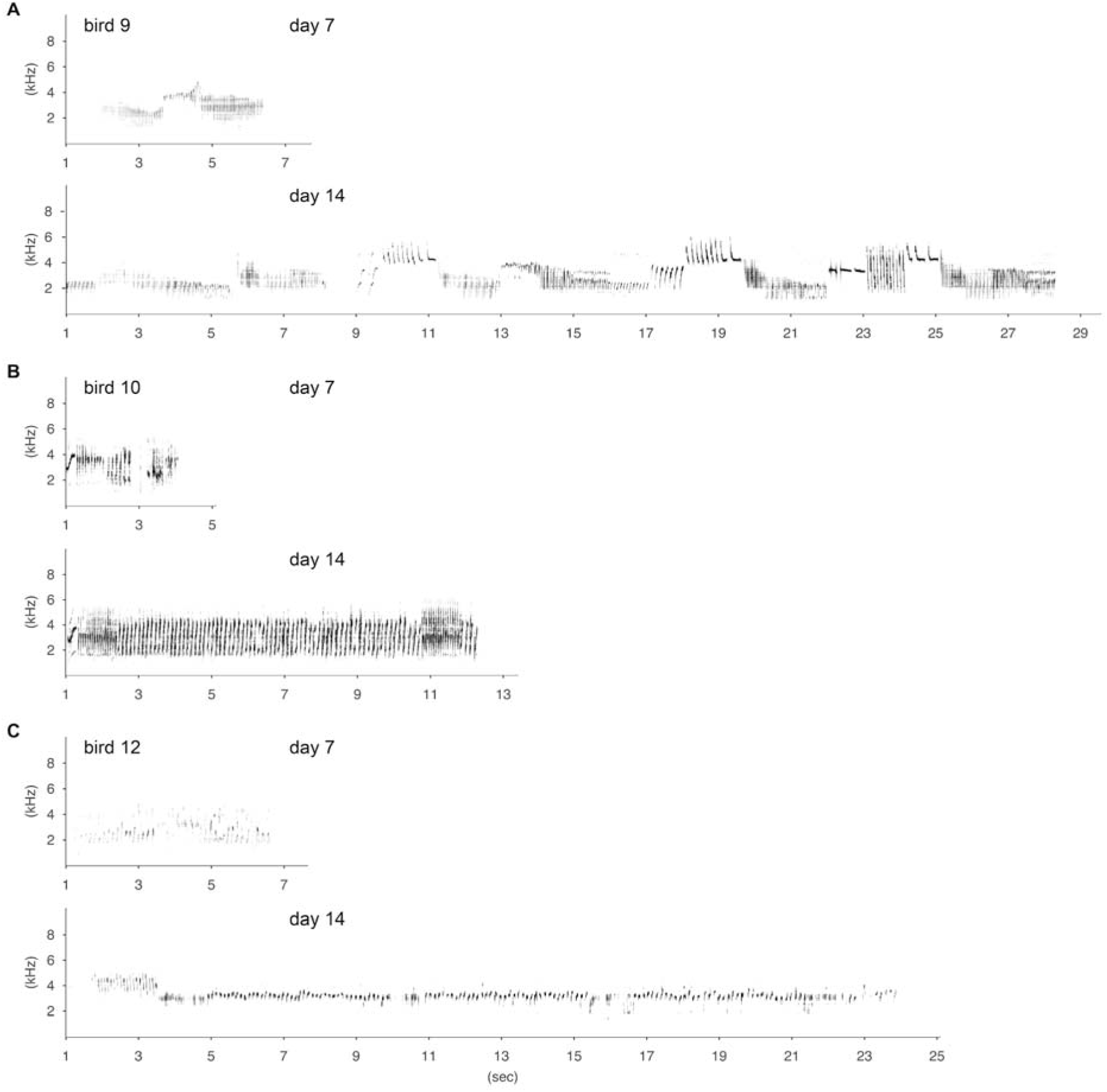
Representative song spectrograms illustrating song development in three T14d birds. Note that the song length, syllable repetitions, and amplitude (the intensity of the spectrogram) of the three birds increased over time. Bird 9 developed the most male-typical song patterns.

**Figure 1 – Figure supplement 2.**
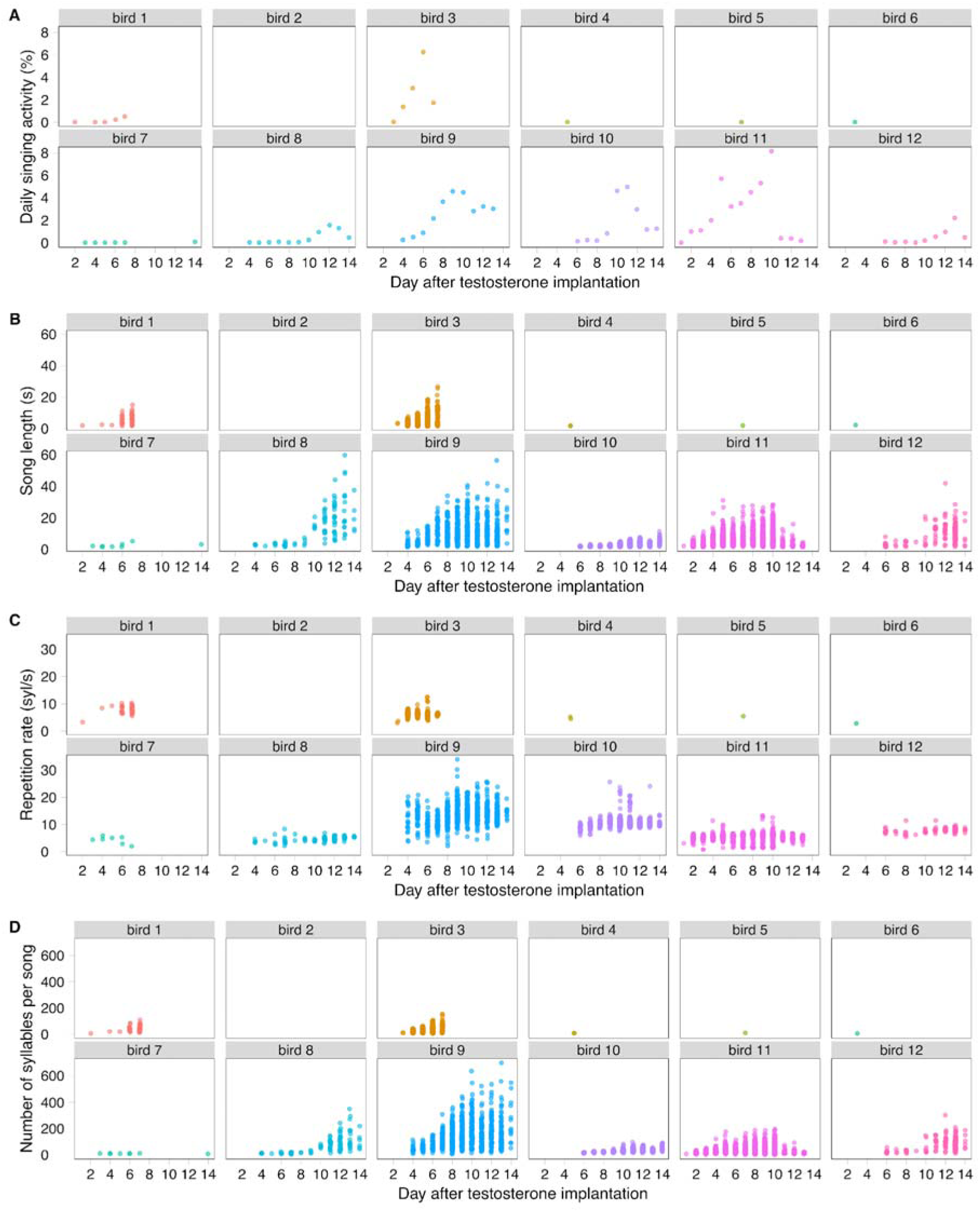
Inter- and intra-individual variability of daily singing activity (A), song length (B), syllable repetition rate (C) and number of syllables per song (D) during song development. Each dot represents a measurement of a day in A and represents a song in B-D. Birds 1 to 6 belong to T7d, and birds 7 to 12 belong to T14d.

**Figure 1 – Figure supplement 3.**
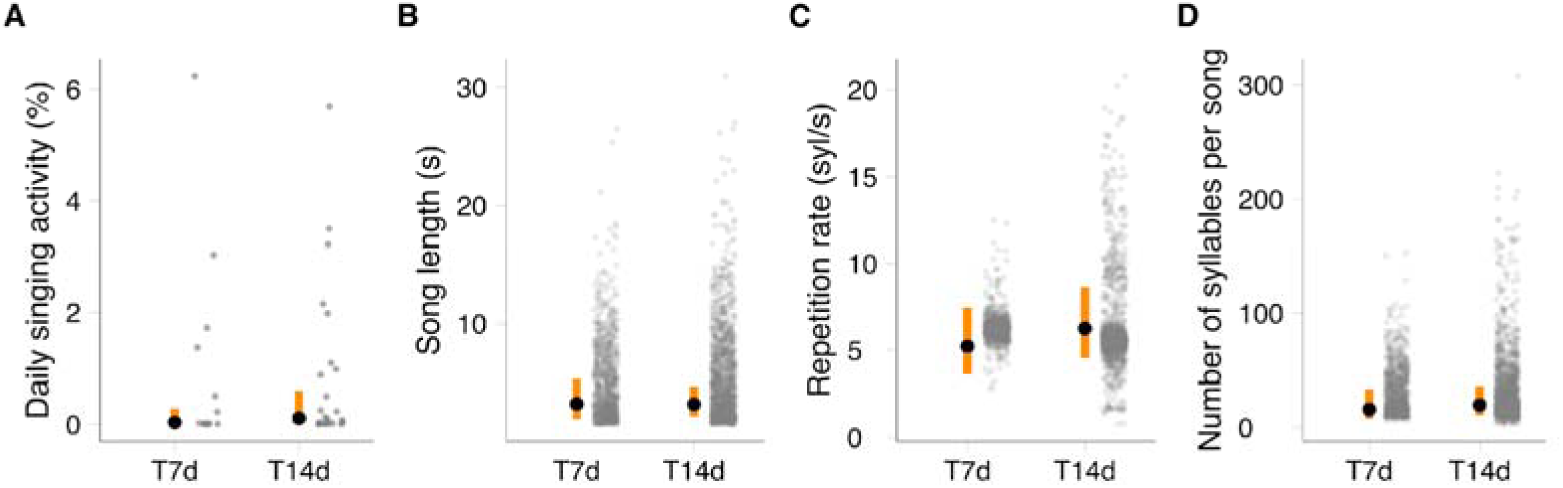
Quantification of song parameters during the first seven days after testosterone implantation in female canaries. **A,** Daily singing activity as measured by the percentage of time spent singing within nine hours per day; **B**, song length in seconds (s); **C**, syllable repetition rate in syllables per second (sys/s); **D**, number of syllables in a song. Each gray dot represents the value per day of a bird in the T7d and T14d groups from day 1 to day 7 after testosterone implantation. The black dots indicate the predicted estimates of the linear mixed-effects models, and the vertical orange bars indicate the 95% credibility intervals (CrI) of the predicted estimates. There are no statistical differences between birds in these two groups, as can be seen from the overlapping 95% CRIs. See also Supplemental Table 2 and Supplemental Table 3 for the linear mixed-effect model estimates and posterior probability.

**Figure 1 – Figure supplement 4.**
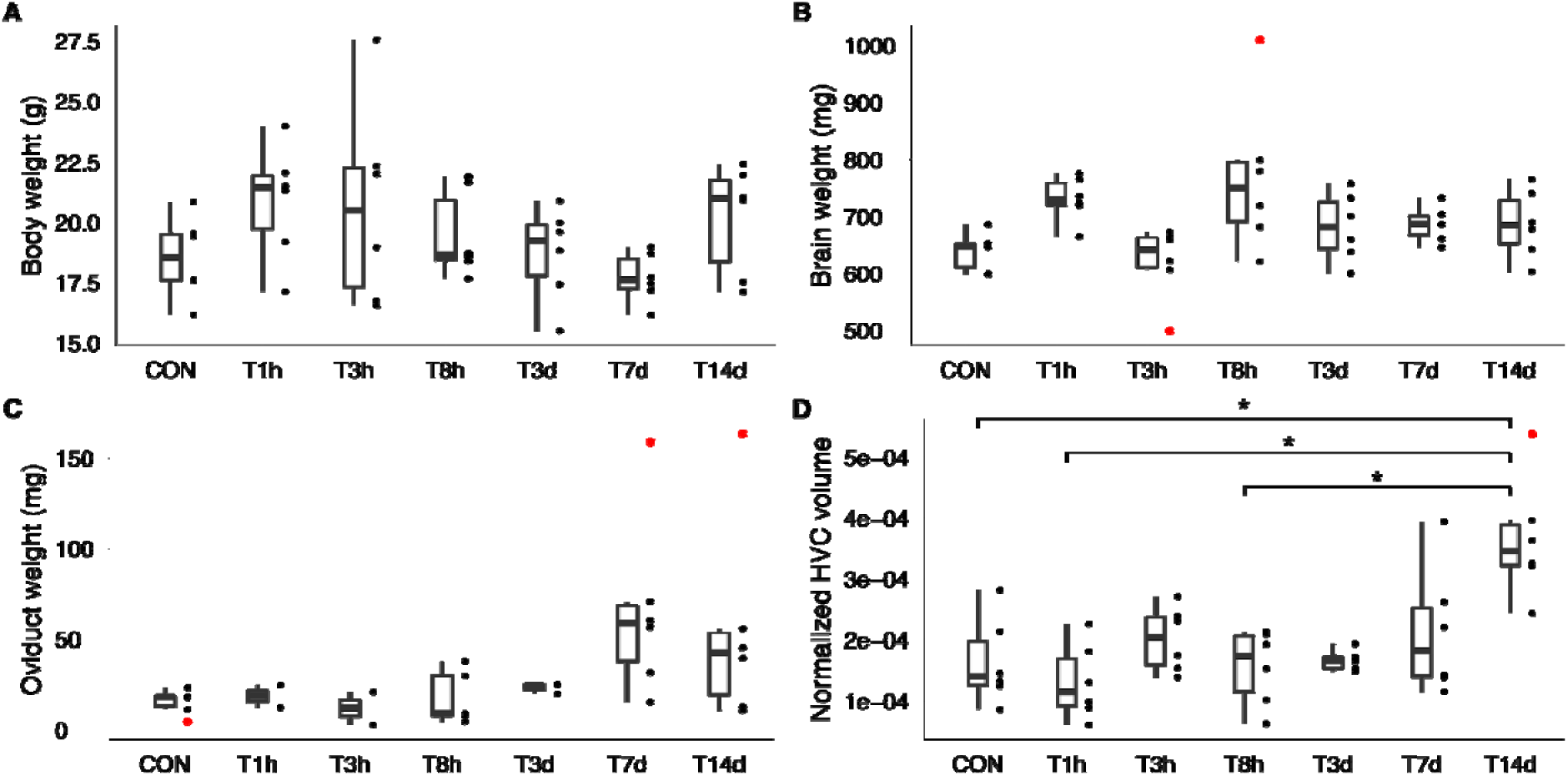
Physiological measurements of testosterone-implanted female canaries. **A**, body weight: No significant differences observed (Kruskal-Wallis rank sum test, χ^2^ = 7.47, df = 6, P-value = 0.28). **B**, brain weight: Significant differences were detected (Kruskal-Wallis rank sum test, χ2 = 15.70, df = 6, P-value = 0.015), but Dunn’s post hoc test with Holm’s adjustment revealed no significant pairwise differences. **C**, oviduct weight: No significant differences observed (Kruskal-Wallis rank sum test, χ2 = 11.58, df = 6, P-value = 0.072). **D**, normalized HVC volume (against brain weight): T14d group significantly different from control (CON), T1h, and T8h groups (Kruskal-Wallis rank sum test, χ2 = 17.32, df = 6, P-value = 0.0082; Holm-adjusted P-values: CON vs. T14d 0.044; T1h vs. T14d 0.0043; T8h vs. T14d 0.046). Boxes indicate 25th/50th/75th percentiles; whiskers show the most extreme values within 1.5 times the inter-quartile range (IQR). Red dots indicate outliers. *: Holm-adjusted P-value <0.05 (Kruskal-Wallis rank sum test followed by Dunn’s post hoc test using the “Holm” method).

**Figure 2 – Figure supplement 1.**
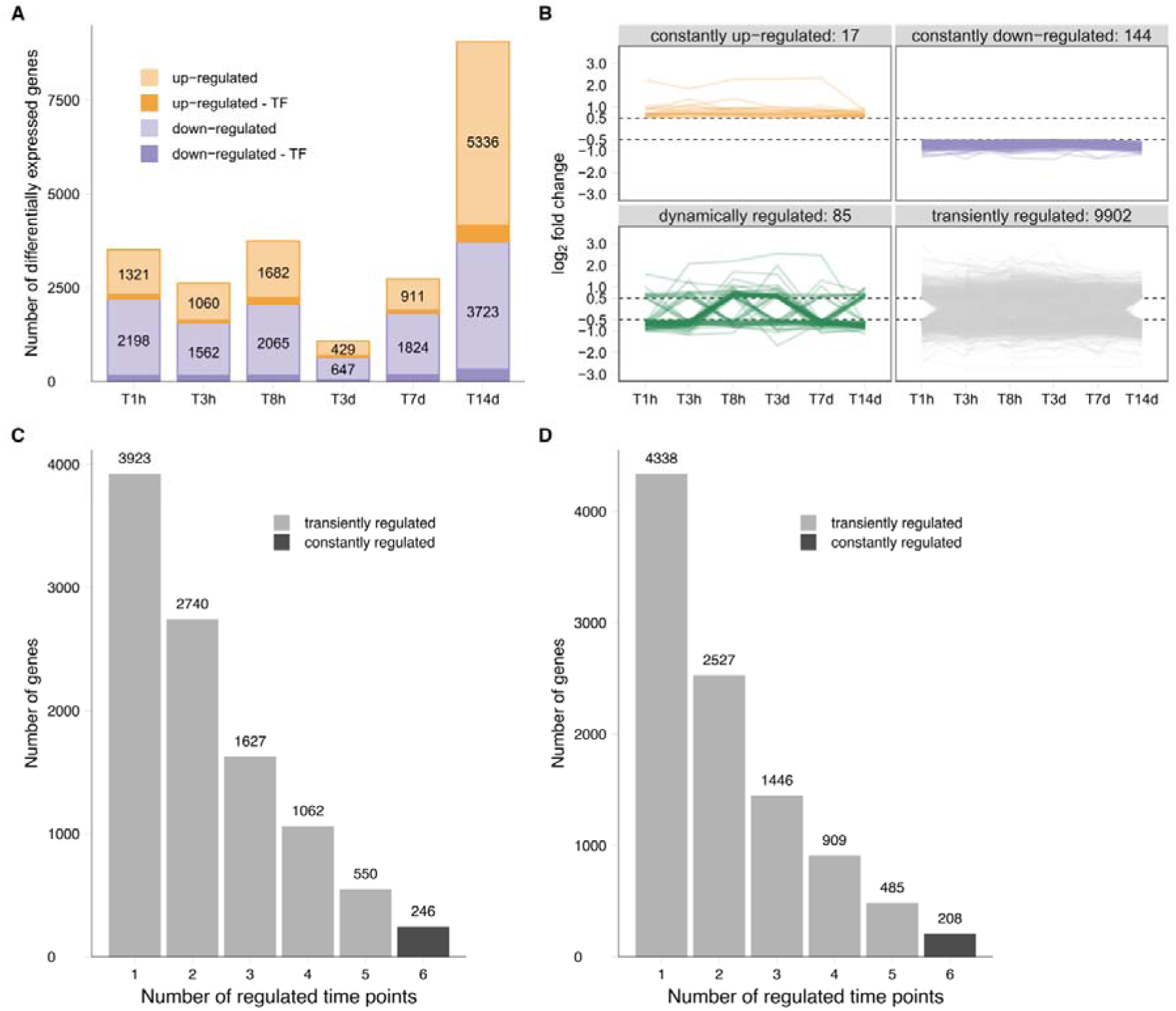
Dissection of transiently regulated genes by the number of differentially regulated time points. **A,** Number of differentially expressed genes in the HVC of testosterone-treated birds compared to controls, including genes with power less than 0.8. Darker shades indicate transcription factors (TF). **B,** Classification of differentially expressed genes, including genes with power less than 0.8, based on temporal regulation patterns: 1) constantly upregulated, 2) constantly downregulated, 3) dynamically regulated, and 4) transiently regulated genes. **C,** Distribution of genes categorized as transiently regulated (light grey) based on the number of time points at which they are differentially expressed. Each bar represents the number of differentially expressed genes at one, two, or more time points, providing a more detailed view of the transiently regulated gene expression patterns. **D,** Similar to C, but excluding genes with power less than 0.8. This provides a more stringent analysis of the transiently regulated genes, focusing on those with higher statistical power.

**Figure 3 – Figure supplement 1.**
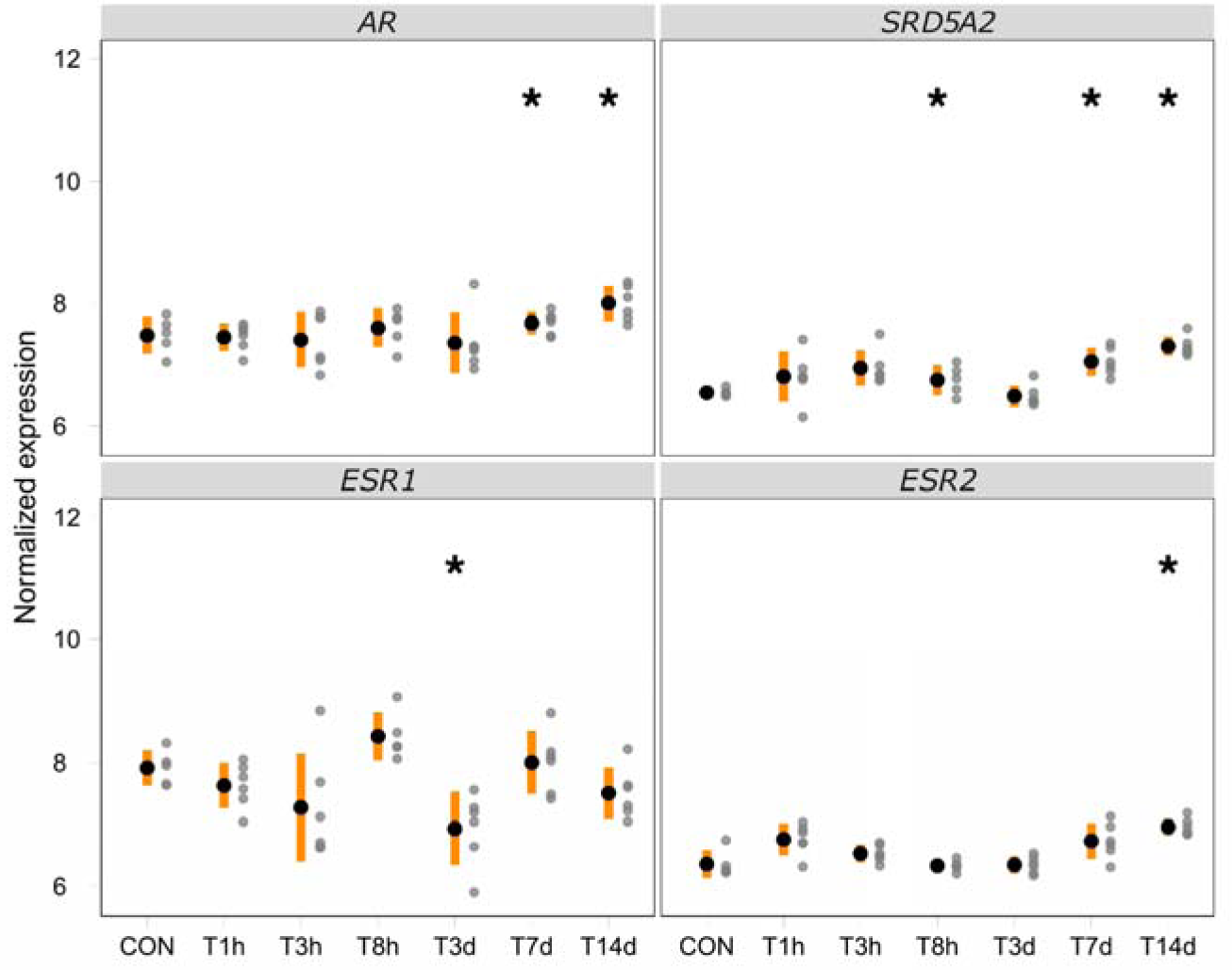
RMA normalized expression levels of selected genes. Each gray dot represents the Robust Multichip Average (RMA)-normalized expression level for an individual bird. Black dots indicate the average expression levels, while vertical orange bars represent the standard deviation of these levels. Asterisks indicate genes that are significantly differentially expressed relative to the control group. The genes shown are *AR*: androgen receptor; *SRD5A2*: 5α-reductase 2; *ESR1*: estrogen receptor α; *ESR2*: estrogen receptor ß.

**Figure 3 – Figure supplement 2.**
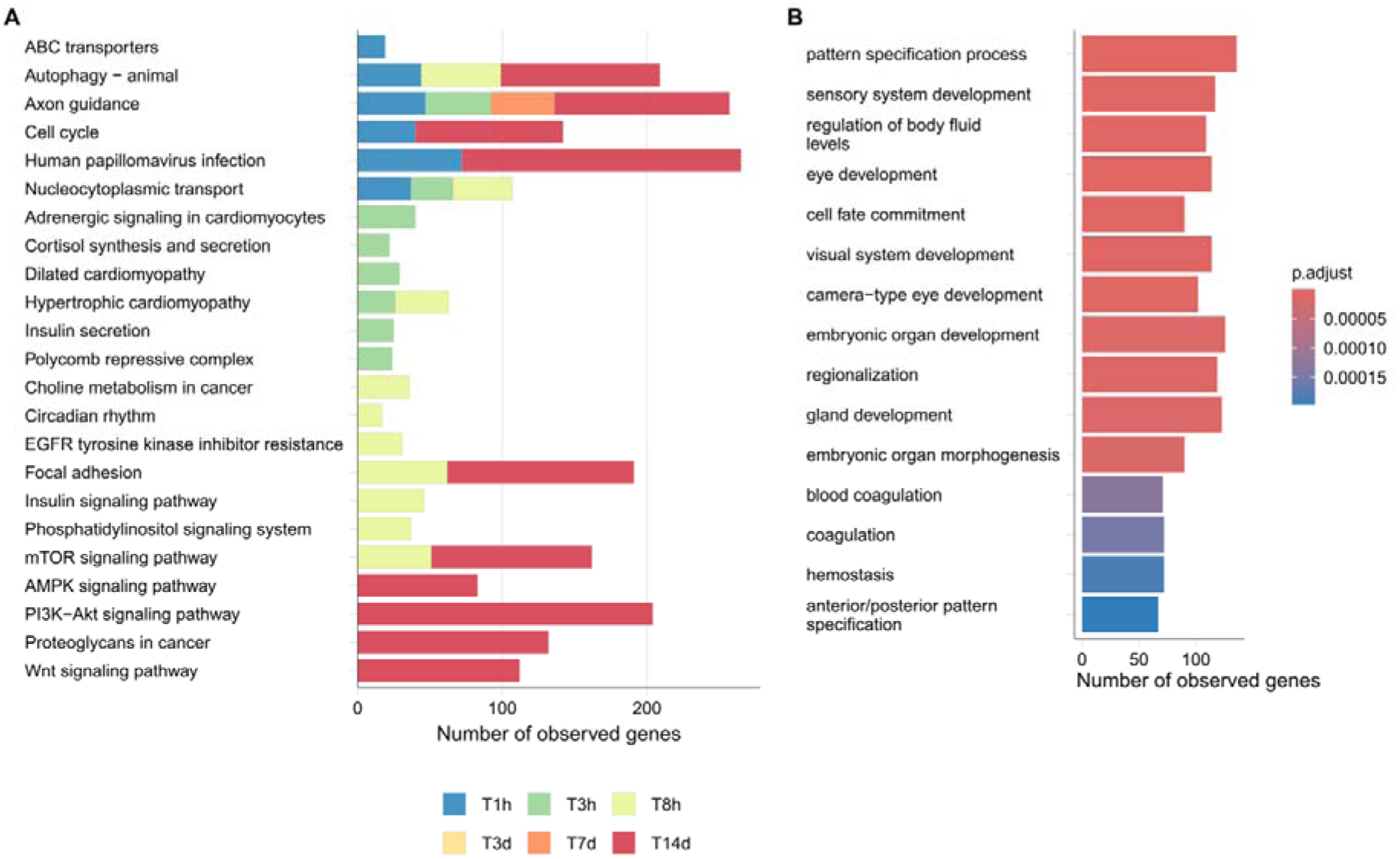
KEGG pathway and GO term enrichment analysis of differentially expressed genes during progressive testosterone action on HVC. **A,** KEGG pathway enrichment analysis of differentially expressed genes at each time point. The number of observed genes (cumulative across time points) is presented in a stacked bar graph. The enriched KEGG pathways (based on Bonferroni-adjusted p-values) are displayed for each time point. Similar to GO term enrichment analysis results, testosterone seems to activate smaller-scale changes, such as focal adhesion, at early time points, while activating system-level pathways like the Hippo signaling pathway, which regulates organ size, at T14d. **B,** GO term enrichment analysis focusing on genes only differentially expressed at T14d, which shows primarily system-level biological processes. The figure shows the ten most probable GO terms based on Bonferroni-adjusted P-values.

## Supplementary Tables and Legends

**Supplementary Table 1.**

**Summary table for measured song parameters (mean ± SD).**

**Supplementary Table 2.**

**Estimates of fixed and random effects of linear mixed effect models.** We present fixed (β) and random (σ2) parameters with their 95% credible intervals (CrI). A statistically meaningful effect of a fixed factor can be assumed if zero is not included within the 95% CrI, and was labeled with *.

**Supplementary Table 3.**

**The posterior probability of a mean difference between compared factors of linear mixed effect models.** Comparisons with statistically meaningful differences, defined by posterior probability > 0.95, were indicated with *.

**Supplementary Table 4.**

**List of differentially expressed genes.** The output of the differential analysis performed using the limma R package. Additional power analysis was performed as described in the methods section “Analysis of differentially expressed genes” of the manuscript.

**Supplementary Table 5.**

**List of constantly regulated genes.**

**Supplementary Table 6.**

**GO-term enrichment analysis of differentially expressed genes of each time point.**

**Supplementary Table 7.**

**KEGG enrichment analysis of differentially expressed genes of each time point.**

**Supplementary Table 8.**

**GO-term enrichment analysis of differentially expressed genes only at T14d.**

**Supplementary Table 9.**

**List of RNAScope probes used in this study.**

## Notes

https://doi.org/10.5061/dryad.5hqbzkh8c

### Competing Interest Statement

The authors have declared no competing interest.

### Summary of Updates

This revised version of the manuscript represents a comprehensive overhaul of our original submission.

## References

[1] Riters LV, Baillien M, Eens M, Pinxten R, Foidart A, Ball GF, et al. Seasonal Variation in Androgen-Metabolizing Enzymes in the Diencephalon and Telencephalon of the Male European Starling (Sturnus vulgaris). J Neuroendocrinol. 2001;13(11):985–997.

[2] Riters LV, Eens M, Pinxten R, Duffy DL, Balthazart J, Ball GF. Seasonal changes in courtship song and the medial preoptic area in male European starlings (Sturnus vulgaris). Horm Behav. 2000;38(4):250–261.

[3] Watts HE. Seasonal regulation of behaviour: what role do hormone receptors play? Proc Biol Sci. 2020;287(1930):20200722.

[4] Ball GF, Balthazart J. Neuroendocrine mechanisms regulating reproductive cycles and reproductive behavior in birds. In: Hormones, Brain and Behavior. Elsevier; 2002:649–XII.

[5] Guh YJ, Tamai TK, Yoshimura T. The underlying mechanisms of vertebrate seasonal reproduction. Proc Jpn Acad Ser B Phys Biol Sci. 2019;95(7):343–357.

[6] Gahr M. Where is the asexual brain? Horm Behav. 2004;46(1):130.

[7] Apfelbeck B, Kiefer S, Mortega KG, Goymann W, Kipper S. Testosterone affects song modulation during simulated territorial intrusions in male black redstarts (Phoenicurus ochruros). PLoS One. 2012;7(12):e52009.

[8] Gahr M. Seasonal hormone fluctuations song structure birds. Coding strategies vertebrate acoustic communication. Published online 2020:163–201.

[9] Alliende J, Giret N, Pidoux L, Del Negro C, Leblois A. Seasonal plasticity of song behavior relies on motor and syntactic variability induced by a basal ganglia–forebrain circuit. Neuroscience. 2017;359:49–68.

[10] Ball GF, Auger CJ, Bernard DJ, Charlier TD, Sartor JJ, Riters LV, et al. Seasonal plasticity in the song control system: multiple brain sites of steroid hormone action and the importance of variation in song behavior. Ann N Y Acad Sci. 2004;1016(1):586–610.

[11] Celec P, Ostatníková D, Hodosy J. On the effects of testosterone on brain behavioral functions. Front Neurosci. 2015;9:12.

[12] Fuentes N, Silveyra P. Estrogen receptor signaling mechanisms. Adv Protein Chem Struct Biol. 2019;116:135–170.

[13] Bao R, Onishi KG, Tolla E, Ebling FJP, Lewis JE, Anderson RL, et al. Genome sequencing and transcriptome analyses of the Siberian hamster hypothalamus identify mechanisms for seasonal energy balance. Proc Natl Acad Sci U S A. 2019;116(26):13116–13121.

[14] Peterson MP, Rosvall KA, Choi JH, Ziegenfus C, Tang H, Colbourne JK, et al. Testosterone affects neural gene expression differently in male and female juncos: a role for hormones in mediating sexual dimorphism and conflict. PLoS One. 2013;8(4):e61784.

[15] Nottebohm F, Nottebohm ME, Crane L. Developmental and seasonal changes in canary song and their relation to changes in the anatomy of song-control nuclei. Behav Neural Biol. 1986;46(3):445–471.

[16] Voigt C, Leitner S. Seasonality in song behaviour revisited: Seasonal and annual variants and invariants in the song of the domesticated canary (Serinus canaria). Horm Behav. 2008;54(3):373–378.

[17] Wild JM. Functional Neuroanatomy of the Sensorimotor Control of Singing. Ann N Y Acad Sci. 2004;1016(1):438–462.

[18] Gahr M. How Hormone-Sensitive Are Bird Songs And What Are The Underlying Mechanisms? Acta Acust United Acust. 2014;100(4):705–718.

[19] Alcami P, Totagera S, Sohnius-Wilhelmi N, Leitner S, Grothe B, Frankl-Vilches C, et al. Extensive GJD2 Expression in the Song Motor Pathway Reveals the Extent of Electrical Synapses in the Songbird Brain. Biology (Basel). 2021;10(11):1099.

[20] Nottebohm F, Stokes TM, Leonard CM. Central control of song in the canary, Serinus canarius. J Comp Neurol. 1976;165(4):457–486.

[21] Gahr M. Localization of androgen receptors and estrogen receptors in the same cells of the songbird brain. Proc Natl Acad Sci U S A. 1990;87:9445–9448.

[22] Johnson F, Bottjer SW. Differential estrogen accumulation among populations of projection neurons in the higher vocal center of male canaries. J Neurobiol. 1995;26(1):87–108.

[23] Hahnloser RHR, Kozhevnikov AA, Fee MS. An ultra-sparse code underliesthe generation of neural sequences in a songbird. Nature. 2002;419(6902):65–70.

[24] Long MA, Fee MS. Using temperature to analyse temporal dynamics in the songbird motor pathway. Nature. 2008;456(7219):189–194.

[25] Frankl-Vilches C, Gahr M. Androgen and estrogen sensitivity of bird song: a comparative view on gene regulatory levels. J Comp Physiol A Neuroethol Sens Neural Behav Physiol. Published online December 6, 2017. doi:10.1007/s00359-017-1236-y

[26] Gahr M. Distribution of sex steroid hormone receptors in the avian brain: Functional implications for neural sex differences and sexual behaviors. Microsc Res Tech. 2001;55(1):1–11.

[27] Louissaint A, Rao S, Leventhal C, Goldman SA. Coordinated Interaction of Neurogenesis and Angiogenesis in the Adult Songbird Brain. Neuron. 2002;34(6):945–960.

[28] Rasika S, Nottebohm F, Alvarez-Buylla A. Testosterone increases the recruitment and/or survival of new high vocal center neurons in adult female canaries. Proc Natl Acad Sci U S A. 1994;91(17):7854–7858.

[29] Nottebohm F. Testosterone triggers growth of brain vocal control nuclei in adult female canaries. Brain Res. 1980;189(2):429–436.

[30] Madison FN, Rouse ML Jr, Balthazart J, Ball GF. Reversing song behavior phenotype: Testosterone driven induction of singing and measures of song quality in adult male and female canaries (Serinus canaria). Gen Comp Endocrinol. 2015;215:61–75.

[31] Hartog TE, Dittrich F, Pieneman AW, Jansen RF, Frankl-Vilches C, Lessmann V. Brain-derived neurotrophic factor signaling in the HVC is required for testosterone-induced song of female canaries. J Neurosci. 2009;29:15511–15519.

[32] Vellema M, Diales Rocha M, Bascones S, Zsebők S, Dreier J, Leitner S, et al. Accelerated redevelopment of vocal skills is preceded by lasting reorganization of the song motor circuitry. Elife. 2019;8. doi:10.7554/eLife.43194

[33] Sartor JJ, Ball GF. Social suppression of song is associated with a reduction in volume of a song-control nucleus in European starlings (Sturnus vulgaris). Behav Neurosci. 2005;119:233–244.

[34] Fusani L, Metzdorf R, Hutchison JB, Gahr M. Aromatase inhibition affects testosterone-induced masculinization of song and the neural song system in female canaries. J Neurobiol. 2003;54:370–379.

[35] Cornez G, Shevchouk OT, Ghorbanpoor S, Ball GF, Cornil CA, Balthazart J. Testosterone stimulates perineuronal nets development around parvalbumin cells in the adult canary brain in parallel with song crystallization. Horm Behav. 2020;119(104643):104643.

[36] Frankl-Vilches C, Kuhl H, Werber M, Klages S, Kerick M, Bakker A, et al. Using the canary genome to decipher the evolution of hormone-sensitive gene regulation in seasonal singing birds. Genome Biol. 2015;16:19.

[37] Poirier K, Lebrun N, Broix L, Tian G, Saillour Y, Boscheron C, et al. Mutations in TUBG1, DYNC1H1, KIF5C and KIF2A cause malformations of cortical development and microcephaly. Nat Genet. 2013;45(6):639–647.

[38] Parcerisas A, Ortega-Gascó A, Pujadas L, Soriano E. The hidden side of NCAM family: NCAM2, a key cytoskeleton organization molecule regulating multiple neural functions. Int J Mol Sci. 2021;22(18):10021.

[39] Rasika S, Alvarez-Buylla A, Nottebohm F. BDNF mediates the effects of testosterone on the survival of new neurons in an adult brain. Neuron. 1999;22(1):53–62.

[40] Bissegger S, Martyniuk CJ, Langlois VS. Transcriptomic profiling in Silurana tropicalis testes exposed to finasteride. Gen Comp Endocrinol. 2014;203:137–145.

[41] Carrier N, Saland SK, Duclot F, He H, Mercer R, Kabbaj M. The Anxiolytic and Antidepressant-like Effects of Testosterone and Estrogen in Gonadectomized Male Rats. Biol Psychiatry. 2015;78(4):259–269.

[42] Cheviron ZA, Swanson DL. Comparative Transcriptomics of Seasonal Phenotypic Flexibility in Two North American Songbirds. Integr Comp Biol. 2017;57(5):1040–1054.

[43] Dopico XC, Evangelou M, Ferreira RC, Guo H, Pekalski ML, Smyth DJ, et al. Widespread seasonal gene expression reveals annual differences in human immunity and physiology. Nat Commun. 2015;6:7000.

[44] Faber-Hammond J, Samanta MP, Whitchurch EA, Manning D, Sisneros JA, Coffin AB. Saccular Transcriptome Profiles of the Seasonal Breeding Plainfin Midshipman Fish (Porichthys notatus), a Teleost with Divergent Sexual Phenotypes. PLoS One. 2015;10(11):e0142814.

[45] Félix AS, Cardoso SD, Roleira A, Oliveira RF. Forebrain Transcriptional Response to Transient Changes in Circulating Androgens in a Cichlid Fish. G3 (Bethesda). 2020;10(6):1971–1982.

[46] Morey JS, Neely MG, Lunardi D, Anderson PE, Schwacke LH, Campbell M, et al. RNA-Seq analysis of seasonal and individual variation in blood transcriptomes of healthy managed bottlenose dolphins. BMC Genomics. 2016;17(1):720.

[47] Quintela T, Marcelino H, Gonçalves I, Patriarca FM, Santos CRA. Gene expression profiling in the hippocampus of orchidectomized rats. J Mol Neurosci. 2015;55(1):198–205.

[48] Sharma A, Das S, Kumar V. Transcriptome-wide changes in testes reveal molecular differences in photoperiod-induced seasonal reproductive life-history states in migratory songbirds. Mol Reprod Dev. 2019;86(8):956–963.

[49] Zhang W, Guo Y, Li J, Huang L, Kazitsa EG, Wu H. Transcriptome analysis reveals the genetic basis underlying the seasonal development of keratinized nuptial spines in Leptobrachium boringii. BMC Genomics. 2016;17(1):978.

[50] Zhao W, Yuan T, Fu Y, Niu D, Chen W, Chen L, et al. Seasonal differences in the transcriptome profile of the Zhedong white goose (Anser cygnoides) pituitary gland. Poult Sci. 2021;100(2):1154–1166.

[51] Gahr M. Delineation of a brain nucleus: comparisons of cytochemical, hodological, and cytoarchitectural views of the song control nucleus HVc of the adult canary. J Comp Neurol. 1990;294:30–36.

[52] Johnson F, Bottjer SW. Hormone-induced changes in identified cell populations of the higher vocal center in male canaries. J Neurobiol. 1993;24(3):400–418.

[53] Colquitt BM, Merullo DP, Konopka G, Roberts TF, Brainard MS. Cellular transcriptomics reveals evolutionary identities of songbird vocal circuits. Science. 2021;371(6530):eabd9704.

[54] Jarvis ED, Nottebohm F. Motor-driven gene expression. Proc Natl Acad Sci U S A. 1997;94(8):4097–4102.

[55] Kilkenny C, Browne WJ, Cuthill IC, Emerson M, Altman DG. Improving bioscience research reporting: the ARRIVE guidelines for reporting animal research. PLoS Biol. 2010;8(6):e1000412.

[56] Griffiths R, Double MC, Orr K, Dawson RJG. A DNA test to sex most birds. Mol Ecol. 1998;7(8):1071–1075.

[57] Ko MC, Van Meir V, Vellema M, Gahr M. Characteristics of song, brain-anatomy and blood androgen levels in spontaneously singing female canaries. Horm Behav. 2020;117:104614.

[58] Tchernichovski O, Nottebohm F, Ho CE, Pesaran B, Mitra PP. A procedure for an automated measurement of song similarity. Anim Behav. 2000;59(6):1167–1176.

[59] Gardner M. White and brown music, fractal curves and one-over-f fluctuations. Published online 1978.

[60] Gisiger T. Scale invariance in biology: coincidence or footprint of a universal mechanism? Biol Rev Camb Philos Soc. 2001;76(2):161–209.

[61] Voss RF, Clarke J. “1/f noise” in music and speech. Nature. 1975;258(5533):317–318.

[62] Wingfield JC, Smith JP, Farner DS. Endocrine Responses of White-Crowned Sparrows to Environmental Stress. Condor. 1982;84(4):399–409.

[63] Goymann W, Möstl E, Gwinner E. Non-invasive methods to measure androgen metabolites in excrements of European stonechats, Saxicola torquata rubicola. Gen Comp Endocrinol. 2002;129(2):80–87.

[64] Rueden CT, Schindelin J, Hiner MC, DeZonia BE, Walter AE, Arena ET, et al. ImageJ2: ImageJ for the next generation of scientific image data. BMC Bioinformatics. 2017;18(1):529.

[65] Schindelin J, Arganda-Carreras I, Frise E, Kaynig V, Longair M, Pietzsch T, et al. Fiji: an open-source platform for biological-image analysis. Nat Methods. 2012;9:676.

[66] Wang F, Flanagan J, Su N, Wang LC, Bui S, Nielson A, et al. RNAscope: A Novel in Situ RNA Analysis Platform for Formalin-Fixed, Paraffin-Embedded Tissues. J Mol Diagn. 2012;14(1):22–29.

[67] Pan W, Lin J, Le CT. How many replicates of arrays are required to detect gene expression changes in microarray experiments? A mixture model approach. Genome Biol. 2002;3(5):research0022.

[68] Ko MC, Frankl-Vilches C, Bakker A, Gahr M. The gene expression profile of the song control nucleus HVC shows sex specificity, hormone responsiveness, and species specificity among songbirds. Front Neurosci. 2021;15:680530.

[69] Dittrich F, Ramenda C, Grillitsch D, Frankl-Vilches C, Ko MC, Hertel M, et al. Regulatory mechanisms of testosterone-stimulated song in the sensorimotor nucleus HVC of female songbirds. BMC Neurosci. 2014;15(1):1–16.

[70] Yates A, Akanni W, Amode MR, Barrell D, Billis K, Carvalho-Silva D, et al. Ensembl 2016. Nucleic Acids Res. 2016;44(D1):D710–D716.

[71] Benson DA, Karsch-Mizrachi I, Lipman DJ, Ostell J, Wheeler DL. GenBank. Nucleic Acids Res. 2005;33(Database Issue):D34–D38.

[72] The UniProt Consortium. UniProt: a hub for protein information. Nucleic Acids Res. 2015;43(D1):D204–D212.

[73] Flicek P, Amode MR, Barrell D, Beal K, Billis K, Brent S, et al. Ensembl 2014. Nucleic Acids Res. 2014;42(D1):D749–D755.

[74] Huang DW, Sherman BT, Lempicki RA. Systematic and integrative analysis of large gene lists using DAVID bioinformatics resources. Nat Protoc. 2008;4(1):44–57.

[75] Huang DW, Sherman BT, Lempicki RA. Bioinformatics enrichment tools: paths toward the comprehensive functional analysis of large gene lists. Nucleic Acids Res. 2009;37(1):1–13.

[76] Edgar R, Domrachev M, Lash AE. Gene Expression Omnibus: NCBI gene expression and hybridization array data repository. Nucleic Acids Res. 2002;30:207–210.

[77] Ritchie ME, Phipson B, Wu D, Hu Y, Law CW, Shi W, et al. limma powers differential expression analyses for RNA-sequencing and microarray studies. Nucleic Acids Res. 2015;43(7):e47.

[78] Klaus B, Reisenauer S. An end to end workflow for differential gene expression using Affymetrix microarrays. F1000Res. 2016;5:1384.

[79] Champely S. Pwr: Basic Functions for Power Analysis.; 2020. https://CRAN.R-project.org/package=pwr. Accessed July 3, 2024

[80] Wu T, Hu E, Xu S, Chen M, Guo P, Dai Z, et al. clusterProfiler 4.0: A universal enrichment tool for interpreting omics data. Innovation (Camb). 2021;2(3):100141.

[81] Wickham H. ggplot2: Elegant Graphics Data Analysis. Springer-Verlag; 2016.

[82] R Core Team. R: A Language and Environment for Statistical Computing. Published online 2024. https://www.R-project.org/

[83] Bates D, Mächler M, Bolker B, Walker S. Fitting linear mixed-effects models Usinglme4. J Stat Softw. 2015;67(1):1–48.

[84] Gelman A, Su YS. arm: Data Analysis Using Regression and Multilevel/Hierarchical Models. Published online 2024. https://CRAN.R-project.org/package=arm

[85] Korner-Nievergelt F, Roth T, Von Felten S, Guélat J, Almasi B, Korner-Nievergelt P. Bayesian Data Analysis Ecology Using Linear Models R, BUGS, Stan. Academic Press; 2015.

